# Signaling events at TMEM doorways provide potential targets for inhibiting breast cancer dissemination

**DOI:** 10.1101/2024.01.08.574676

**Authors:** Chinmay R. Surve, Camille L. Duran, Xianjun Ye, Xiaoming Chen, Yu Lin, Allison S. Harney, Yarong Wang, Ved P. Sharma, E. Richard Stanley, Dianne Cox, John C. McAuliffe, David Entenberg, Maja H. Oktay, John S. Condeelis

## Abstract

Tumor cell intravasation is essential for metastatic dissemination, but its exact mechanism is incompletely understood. We have previously shown that in breast cancer, the direct and stable association of a tumor cell expressing Mena, a Tie2^hi^/VEGF^hi^ macrophage, and a vascular endothelial cell, creates an intravasation portal, called a “tumor microenvironment of metastasis” (TMEM) doorway, for tumor cell intravasation, leading to dissemination to distant sites. The density of TMEM doorways, also called TMEM doorway score, is a clinically validated prognostic marker of distant metastasis in breast cancer patients. Although we know that tumor cells utilize TMEM doorway-associated transient vascular openings to intravasate, the precise signaling mechanisms involved in TMEM doorway function are only partially understood. Using two mouse models of breast cancer and an *in vitro* assay of intravasation, we report that CSF-1 secreted by the TMEM doorway tumor cell stimulates local secretion of VEGF-A from the Tie2^hi^ TMEM doorway macrophage, leading to the dissociation of endothelial junctions between TMEM doorway associated endothelial cells, supporting tumor cell intravasation. Acute blockade of CSF-1R signaling decreases macrophage VEGF-A secretion as well as TMEM doorway-associated vascular opening, tumor cell trans-endothelial migration, and dissemination. These new insights into signaling events regulating TMEM doorway function should be explored further as treatment strategies for metastatic disease.

## Introduction

Metastasis is a complex, multi-step and multi-directional process that begins with the dissemination of tumor cells from the primary tumor to distant sites and escalates by further re-dissemination of cancer cells from metastatic foci (1–4). The re-dissemination process leads to an exponential increase in tumor burden and, eventually, patient demise. Tumor cell intravasation into the vasculature creates circulating tumor cells (CTCs) and is an essential step in the metastatic cascade, but its exact mechanism is not completely understood. A large body of evidence indicates that tumor cell dissemination, including tumor cell intravasation, is orchestrated by the interaction of tumor cells with the tumor microenvironment (5, 6). In particular, tumor-associated macrophages interact with tumor cells and promote directed cell migration (streaming) towards blood vessels (7–9). In addition, macrophages form a multi-cellular complex with endothelial and tumor cells on the surface of blood vessels to mediate transient vascular opening and tumor cell intravasation. These cell complexes are called TMEM (Tumor MicroEnvironment of Metastasis) doorways (10, 11). Each TMEM doorway is composed of one tumor cell expressing Mena, one Tie2^hi^/VEGF-A^hi^ macrophage, and a vascular endothelial cell, all in direct and stable physical contact (10, 12, 13).

TMEM doorway-associated vascular opening (TAVO) was first discovered using high-resolution intravital imaging as localized and transient bursts of intravascular contrast agents into the extravascular space that could be accompanied by concurrent intravasation of cancer cells (10). Further analyses revealed in mouse models of breast cancer, TAVO, and its associated cancer cell intravasation, occurs exclusively at TMEM doorways (10). Importantly, TMEM doorway score in primary breast tumors has been clinically validated as a prognostic marker of distant metastatic recurrence in breast cancer patients with estrogen receptor positive HER2 negative (ER^+^/HER2^-^) disease, independent of other clinical prognostics (12–14). Additionally, when comparing residual ER^+^/HER2^-^ tumors from breast cancer patients after completion of pre-operative chemotherapy, Black patients were found to have a higher TMEM doorway score than white patients (15). These findings potentially provide an explanation for the persistent racial disparities in ER^+^/HER2^-^ breast cancer outcomes that are not fully explained by disparities in social determinants of health, including access to care or treatment (12, 15–17). TMEM doorway score has also been found to increase during neo-adjuvant chemotherapy, suggesting that TMEM doorway score, as measured using TMEM doorway activity MRI (17), may be a useful companion diagnostic for the prediction of benefit from certain types of treatment. Given that Black compared to white patients have a higher TMEM doorway density in their residual disease after chemotherapy, using TMEM doorway activity MRI may potentially help diminish racial disparity in breast cancer outcome (18, 19).

In humans and mice, TMEM doorways are found in pre-invasive and invasive ductal breast carcinoma, as well as in metastatic foci in lymph nodes and lungs (18–24). This suggests that the TMEM doorway-mediated mechanism of cancer cell dissemination occurs not only at the primary tumor site, but also at metastatic sites, and may contribute to the multi-directional cancer spread that perpetuates metastatic dissemination even after removal of the primary tumor (1–4, 18). Thus, understanding the molecular mechanisms of TMEM doorway function may help develop therapeutic targets to slow distant metastases and improve patient survival.

Inhibiting TAVO prevents the formation of CTCs and metastases (10). Although we know that TMEM doorways induce localized vascular opening through the secretion of vascular endothelial growth factor-A (VEGF-A) from the TMEM doorway macrophage (10), the precise signaling mechanisms involved in TMEM doorway function during intravasation have not been elucidated. The work described here provides a greater understanding of the signaling events that regulate these dynamic, multi-cellular interactions. In particular, we report the identification of the signaling events involved in TAVO and TMEM doorway-mediated intravasation, which can serve as potential new prognostic markers and therapeutic targets for suppressing CTC dissemination and breast cancer metastasis.

## Materials and Methods

### Cell culture

MDA-MB-231 cells were cultured in Dulbecco’s Modified Eagle Medium (DMEM) (cat# SH30243, Hyclone, GE Healthcare Life Sciences, Logan, UT, USA) supplemented with 10% fetal bovine serum (FBS) (cat# S11550, Atlanta Biologicals, Flowery Branch, GA, USA). BAC1.2F5 macrophages (67) and bone marrow derived macrophages (BMMs) were cultured in Minimum Essential Medium, Alpha (α-MEM) (cat# 15-012-CV, Corning, Tewksbury, MA, USA) supplemented with 10% FBS (cat# 100-106, Gemini Bio-Products, Sacramento, CA, USA) and 36 µg/mL CSF-1 (3000 units/mL, a gift from Dr. E. Richard Stanley). Immortalized BMMs derived from *Csf1r^+/+^* and *Csf1r^-/-^* mice have been previously described (33), and were cultured in α-MEM supplemented with 10% FBS and GM-CSF. Human Umbilical Vein Endothelial Cells (HUVECs) were cultured in complete EGM-2 (cat# CC-3162, Lonza, Allendale, NJ, USA) and were not used beyond passage five for any experiments.

### Mice

All studies involving mice were conducted in accordance with the National Institutes of Health regulations concerning the care and use of experimental animals. The procedures were approved by the Albert Einstein College of Medicine Institute for Animal Care and Use Committee. Transgenic mice expressing the Polyoma Middle T (PyMT) antigen under the control of the mammary tumor virus long terminal repeat (MMTV-LTR) (68) were bred in house.

### Inhibitors

Inhibitors were administered based on established doses of efficacy found in previous studies. Animals were treated with 2.5 μg of anti-CSF-1R neutralizing antibody (cat# NBP1-43363, clone AFS98, Novus Biologicals, Centennial, CO, USA), or antibody isotype control (cat# 554682, IgG K isotype control, BD Pharmingen, Franklin Lakes, NJ, USA) was administered by tail vein *i.v.* four hours before termination of the experiment as previously described (27). For *in vitro* studies, GW2580 a small molecule inhibitor of CSF-1R (cat# G-5903, LC Laboratories, Woburn, MA, USA) was dissolved in DMSO was used at a concentration of 100 nM, and anti-CSF-1R neutralizing antibody or isotype control (clone AFS98, Novus Biologicals) was used at a concentration of 50 ng/mL

### Immunofluorescence labeling of tumor vasculature and extravasation with 155 kDa dextran-TMR

Labeling flowing vasculature and sites of permeability was performed as previously described (10). Briefly, to quantify extravasation, high molecular weight 155 kDa TMR-dextran (cat# T1287, Sigma-Aldrich, Burlington, MA, USA) diluted in PBS was administered by tail vein *i.v.* one hour before the termination of the experiments. Anti-mouse CD31-biotin (cat# 13-0311-82, Thermo Fisher, Waltham, MA, USA) was administered by tail vein *i.v.* ten minutes before the end of the experiment to label flowing blood vessels. At time of sacrifice tumors were removed and fixed for 48 hours in 10% formalin in a volume ratio of tumor to formalin of 1:7 and made into paraffin blocks. Paraffin blocks of tumors were cut into 5 µm sections and immunofluorescence staining was performed. TMR-Dextran is visualized using rabbit anti-TMR (A-6397; Life Technologies, Carlsbad, CA, USA).

### Immunofluorescence staining of tissue

Tumor sections were dewaxed in acetone and rehydrated in alcohol followed by water. Antigen retrieval was performed with a citrate solution at pH 5.5 in a humidified chamber. The slides were washed with PBS-T (0.1% Tween-20) and blocked with a blocking solution (2% BSA, 10% FBS in PBS). One 5 μm section from each tumor was stained for hematoxylin and eosin (H&E) and one for TMEM doorway. TMEM doorway stain is a triple immuno-stain for predicting metastatic risk, in which three antibodies are applied sequentially and developed separately with different chromogens on a Dako Autostainer (13, 14, 32). The following primary antibodies were used for immunostaining of TMEM doorways: anti-Mena (cat# NBP1-87914, Novus Biologicals), anti-endomucin (cat# sc-65495, Santa Cruz Biotechnology, Dallas, TX, USA), and anti-CD68 (cat# MCA1957, clone FA-11, Serotec, Kidlington, UK), or anti-Iba1 (anti-Iba1 (cat# 019-19741, FUJIFILM Wako Chemicals, Richmond, VA). The sequential tissue sections were stained with different combinations of: anti-endomucin (cat# sc-65495, Santa Cruz Biotechnology), anti-TMR (to visualize TMR-dextran, cat# A-6397, Thermo Fisher Scientific), anti-ZO-1 (cat# MABT11, clone R40.76, Millipore Sigma), anti-CD31 (cat# 77699; Cell Signaling Technology, Danvers, MA, USA), Red fluorochrome(635)-conjugated anti-Iba1 (cat# 013-26471, FUJIFILM Wako Chemicals), or anti-Iba1 (cat# 019-19741, FUJIFILM Wako Chemicals), anti-VEGF-A (cat# sc-152, clone A-20, Santa Cruz Biotechnology) or anti-VEGF-A (cat#512809, clone 2G11-2A05, Biolegend, San Diego, CA, USA), anti-CSF-1R (cat# sc-692, clone C-20, Santa Cruz Biotechnology), AlexaFluor555-conjugated anti-CSF-1 (cat# bs-4910R-A555, Bioss Inc, Woburn, MA, USA), anti-CD68 (cat# MCA1957, clone FA-11, Serotec), or AlexaFluor647-conjugated CD68 (cat#51-0689-42, eBioscience, San Diego, CA, USA), and anti-MRC1/CD206 (cat# AF2535, R&D Systems, Minneapolis, MN, USA). Sections were washed with PBS-T and the primary antibodies were detected with AlexaFluor 488, 555 or 647 conjugated secondary antibodies targeting the primary antibody species (Invitrogen, Eugene, OR, USA), and nuclei were stained with 4, 6-diamidino-2-phenylindole (DAPI). All fluorescently labeled samples were mounted with Prolong Diamond antifade reagent (cat# P36961, Invitrogen) and imaged with a PerkinElmer Pannoramic 250 Flash II digital whole-slide scanner using a 20x 0.8NA Plan-Apochromat objective (PerkinElmer, Hopkinton, MA, USA). Images of individual fields of view were imported into ImageJ or VisioPharm for analysis. TMEM doorways were identified as previously described (10, 13, 14). Vascular junctions and extravascular dextran were quantified as previously described (10, 30). Briefly, the CD31 channel (blood vessel), dextran and vascular junction (ZO-1) were each thresholded to just above background based upon intensity and by comparison to a slide stained only with secondary antibodies. A binary mask was created for the CD31 positive staining (blood vessel mask) and for the dextran positive staining (dextran mask). The extravascular dextran area was isolated by subtracting the blood vessel mask from the dextran mask. The remaining extravascular dextran area and blood vessel area were then measured.

### Extravascular dextran, VEGF-A, and intracellular CSF-1 quantification in TMEM doorways

To identify active TMEM doorways, we utilized an improved and automated method of a prior TMEM activity assay that takes into account the presence of high-molecular weight dextran that leaks into the tissue, as a result of TMEM doorway associated vascular opening (TAVO) (10, 30, 31). The algorithmic improvement of this assay involves the automated measurement of the entire tissue section of the tumor as a single region of interest (ROI), instead of selecting a definite number of high power fields as ROIs, thus providing a more accurate representation of the entire tumor, minimizing operator-dependent biases, and significantly increasing the logistic capacity of the analysis.

In short, serial tissue sections were cut and stained for TMR-Dextran, CSF-1, and Endomucin by IF (tissue section 1) and for TMEM doorways by staining for Iba1, Endomucin, and panMena by IHC (tissue section 2). The sections were then imaged with a digital whole slide scanner and aligned to the single cell level using the TissueAlign module in Vis (Visiopharm, Hoersholm, Denmark). IF images were thresholded above background, creating binary masks for endomucin (blood vessel) and dextran signals. These masks were then superimposed so as to be able to differentiate between intravascular and extravascular dextran signal. In the IHC section, TMEM doorways were identified using previously published criteria (13, 14). In this manner, the amount of extravascular dextran around TMEM doorways could be quantified. For quantification of intracellular CSF-1, the TMEM doorway associated tumor cell was identified using the Mena stain in IHC section, within the TMEM doorway circle, and a mask was created. This mask was then overlaid onto the aligned IF section and the CSF-1 was then quantified within this mask.

### siRNA knockdown

MDA-MB-231 cells were transfected with Stealth RNAi siRNAs (siRNA IDs: HSS102355, HSS102356, HSS175321, Thermo Fisher Scientific) directed against CSF-1 or negative control (cat#12935300, Thermo Fisher Scientific) at a concentration of 200 pmols using Lipofectamine 2000 reagent (cat# 11668027, Invitrogen).

### Immunofluorescence staining of co-cultured cells

Prior to the start of the experiment, macrophages were labeled with CellTracker Green^TM^ dye and tumor cells were labelled with CellTracker^TM^ Red dye, according to manufacturer’s instructions (cat# C7025, C34552, Invitrogen). 1×10^5^ macrophages were cultured with or without 1×10^5^ tumor cells in α-MEM media (supplemented with 0.5% FBS and 300 units/mL of CSF-1) and with either CSF-1R blocking antibody or isotype control antibody at 50 ng/mL (cat# NBP1-43363, clone AFS98, Novus Biologicals; IgG K isotype control cat# 554682, BD Pharmingen). After 24 hours incubation, the experiments were stopped by washing the cells with ice-cold PBS followed by fixing them for twenty minutes in 4% PFA, and permeablizing in 0.1% Triton X-100 in PBS. The cells were stained for VEGF-A using antibodies against mouse VEGF-A (cat# 512809, clone 2G11-2A05, Biolegend) and AlexaFluor647-conjugated secondary antibodies. The intensity of VEGF-A staining in the macrophages was measured by first outlining the macrophages in the green (488 nm) channel in ImageJ/Fiji (69) and applying that outline to the far-red (647 nm) channel to measure the VEGF-A signal within the macrophage, excluding any VEGF-A signal from the tumor cells.

### Quantitative real-time polymerase chain reaction (qPCR)

To quantify gene expression in MDA-MB-231 tumor cells transfected with CSF-1 siRNA or control siRNA, tumor cells were grown under normal culture conditions. In co-culture experiments between tumor cells and macrophages, 1×10^6^ macrophages were cultured alone or co-cultured with 1×10^6^ MDA-MB-231 tumor cells for 24 hours in α-MEM media with 0.5% FBS and 300 units/mL of CSF-1 (one-tenth of CSF-1 added to complete media). For the co-culture assay, species specific primers were used in the qPCR.

Total RNA was isolated from cells by using the RNeasy Plus Mini Kit (cat# 74134, Qiagen, Germantown, MD, USA) and cDNA was synthesized and amplified from 1 μg total RNA using the using SuperScript IV VILO Master Mix with ezDNase Enzyme (cat# 11766050, Thermo Fisher Scientific) per manufacturer’s protocol. The qPCR was performed with SYBR Green PCR Master Mix (cat# 4367659, Thermo Fisher Scientific) using a QuantStudio 3 real-time PCR instrument (applied biosystems, Thermo Fisher Scientific). The following primers were used: mouse GAPDH 5’- CTCATGACCACAGTCCATGC -3’, 5’- CACATTGGGGGTAGGAACAC -3’; mouse VEGF-A 5’- AGCAGAAGTCCCATGAAGTGA -3’, 5’- ATGTCCACCAGGGTCTCAAT -3’; human GAPDH 5’- CTCCTGTTCGACAGTCAGCC -3’, 5’- ACCAAATCCGTTGACTCCGAC -3’; human β-Actin 5’- CTTCGCGGGCGACGATGC -3’, 5’- CGTACATGGCTGGGGTGTTG -3’; human CSF-1 5’- CCTCCCACGACATGGCT -3’, 5’- GAGACTGCAGGTGTCCACTC -3’. The mean cycle threshold (Ct) values were then used to analyze relative expression. Analysis was performed using comparative-CT method (2^−ΔCT^ method) and all Ct values were normalized to GAPDH. Each reaction was performed in triplicate.

### ELISA

In macrophage-tumor cell co-culture experiments, 1×10^6^ macrophages were cultured alone or co-cultured with 1×10^6^ MDA-MB-231 tumor cells for up to 48 hours in α-MEM media with 0.5% FBS and 300 units/mL of CSF-1 (one-tenth of CSF-1 added to complete media). Experiments using immortalized *Csf1r^+/+^* or *Csf1r^-/-^* BMMs used α-MEM media with 0.5% FBS and 2 ng/mL GM-CSF. In mono-culture macrophage ELISA experiments where tumor cell conditioned media was added, conditioned media was collected from tumor cells incubated in α-MEM with 0.5% FBS and 300 units/mL of CSF-1 for 24 hours. In tumor cell mono-culture ELISA experiments, 1×10^6^ tumor cells were seeded and incubated for 24 hours in serum-free DMEM medium.

In all ELISA experiments, blocking antibodies, isotype controls, inhibitors and controls were added at indicated concentrations in “Inhibitors” section, at time zero. Media conditioned by the cultured cells were collected at the indicated time points and stored at -80^0^C until use. ELISAs were performed per the manufacturer’s recommendations using the Mouse VEGF-A DuoSet ELISA kit (cat# DY493, R&D Systems) or the Human M-CSF DuoSet ELISA kit (cat# DY216, R&D Systems). The concentration of protein secreted was interpolated from the standard curve measurements.

### Transendothelial Migration Assay (iTEM assay)

The iTEM assay was performed as previously described (34, 70, 71) and briefly described here. The transwell (8 µm pore size; cat# 353097, Corning, Corning, NY, USA) was prepared so that tumor cell transendothelial migration was in the intravasation direction found *in vivo* (from subluminal side to luminal side of the endothelium). To prepare the endothelial monolayer, the underside of each transwell was coated with 50 µL of Matrigel (2.5 µg/mL; cat# 354230, Corning). Approximately 1×10^5^ HUVECs were plated on the Matrigel coated underside of the transwell. Transwells were then flipped into a 24-well plate containing 200 µL of EGM-2 and monolayers were formed over a 48 hour period. The integrity of the endothelium was measured using low molecular weight dextran as described previously (71). Macrophages and tumor cells were labelled with cell tracker dyes (cat# C7025, C34552, Invitrogen) before the experiment. Then, 1.5×10^4^ tumor cells and 6×10^4^ macrophages were added to the upper chamber of the transwell in 200 µL of DMEM without serum while the bottom chamber contained EGM-2 supplemented with 36 µg/mL of CSF-1. After 18 hours, the transwells were fixed and stained for ZO-1 (cat# 402200, Invitrogen) as previously described. Transwells were imaged using a Leica SP5 confocal microscope using a 60x 1.4NA objective and processed using ImageJ/Fiji (69). Tumor cell transendothelial migration quantification was performed by counting the number of tumor cells that had crossed the intact endothelium (intact monolayers confirmed by ZO-1 staining for tight junctions) within the same field of view (60x, 10 random fields) and represented as normalized values from at least three independent experiments.

### Statistical analysis

Individual animals in each cohort are presented as individual points on a dot plot. A horizontal line indicates the mean value and the error bars represent the standard error of the mean. Statistical significance was determined using an unpaired Student’s *t*-test, one-way ANOVA, or two-way ANOVA, with Tukey’s multiple comparisons test, as indicated, using GraphPad Prism (version 10; Graph Pad Software, La Jolla, CA). Data sets were checked for normality (D’Agostino & Pearson omnibus normality test or Shapiro-Wilk normality test) and unequal variance using GraphPad Prism. Welch’s correction was applied to *t*-tests as needed. *P* values less than 0.05 were deemed significant. For *in vitro* experiments, results are representative of at least three independent experiments.

## Results

### CSF-1 levels are elevated in tumor cells at active TMEM doorways

Macrophage-secreted VEGF-A is critical for regulation of TMEM doorway activity (10). VEGF-A secreted by macrophages is regulated by CSF-1, also known as M-CSF (25, 26). Since tumor cells are known to secrete CSF-1 (27–29), we investigated if TMEM doorway tumor cells express CSF-1, and if this expression is associated with TMEM doorway vascular opening (10). We first determined whether CSF-1 levels are increased in TMEM doorway tumor cells at active TMEM doorways by multiplex staining of breast tumor tissue sections obtained from the *polyoma* middle T antigen (PyMT) mice injected *i.v.* with high molecular weight dextran (155 kDa, green) 1 hour before sacrifice. In these samples, if there has been a TMEM doorway associated vascular opening (TAVO) (which would indicate an active TMEM doorway), the *i.v.* injected dextran would extravasate into the interstitium and be detectable extravascularly in the tissue near a TMEM doorway **(Figure 1A)**. We used a modified approach of our previously published protocol for detection of active TMEM doorways (30). We stained two sequential tumor sections – one using IHC for TMEM doorways (Iba1, Endomucin, Mena) **(Figure 1B)** and the second using IF for CSF-1, dextran, and endothelial cells (endomucin) **(Figure 1C)** and aligned the images of stained sections to the single cell level. Next, we identified which TMEM doorways were active by examining for the presence of high molecular weight dextran (155 kDa, green) in the extravascular space near the TMEM doorway, as described previously (10, 31). We then visualized and quantified the expression of CSF-1 (red) in TMEM doorway tumor cells of both active and inactive doorways **(Figure 1C)**. We observed that the fluorescence intensity of intracellular CSF-1 in TMEM doorway tumor cells was higher in active TMEM doorways **(Figure 1C, lower panel)** compared to inactive TMEM doorways **(Figure 1C, upper panel)**. This is quantified in **Figure 1D**. These results indicate that TMEM doorway tumor cells may play a significant role in regulating TMEM doorway function and activity by locally producing CSF-1.

**Figure 1.**
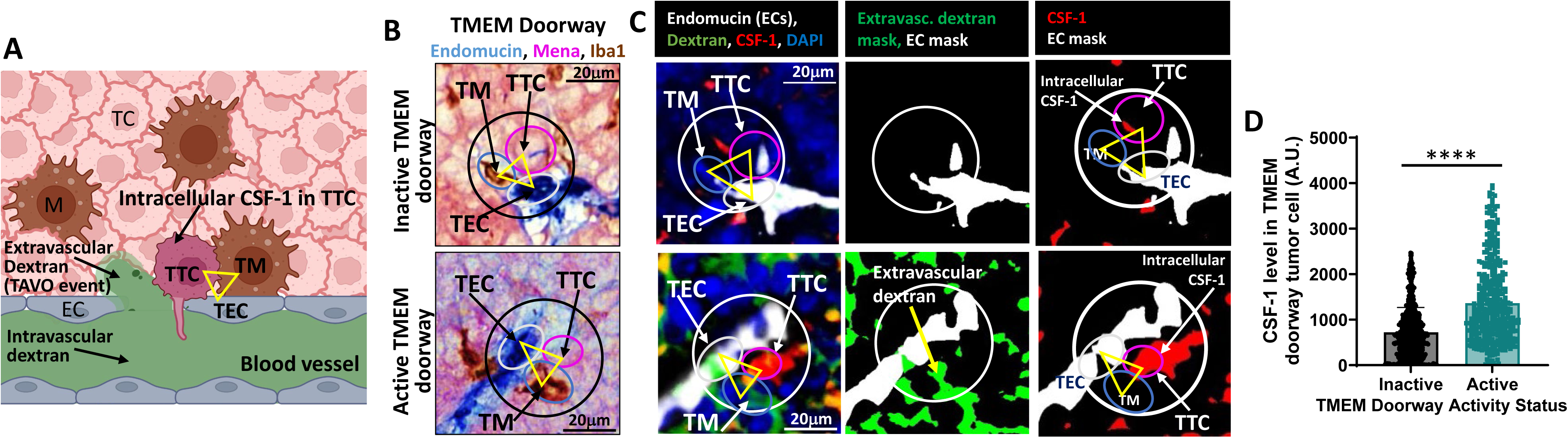
TMEM doorway tumor cells show increased CSF-1 levels at active TMEM doorways. **A)** Cartoon denoting the identification of parameters used for analysis of CSF-1 levels in TMEM doorway tumor cells (TTCs) and TMEM doorway function. The three cells composing the TMEM doorway are indicated by the yellow triangle connecting the TMEM doorway macrophage (TM), the TMEM doorway endothelial cell (TEC) and the TTC. Intravascular dextran is in the lumen of blood vessel, extravascular dextran is outside the lumen of blood vessel, and CSF-1 is directly measured in the TTC. Figure created with BioRender.com. **B)** Panel shows inactive and active TMEM doorways, visualized by immunohistochemistry (IHC) staining for Mena, Iba-1, and endomucin. The three cells of the TMEM doorway (contained in black circle, with the three cells forming the TMEM doorway indicated with the yellow triangle) are the TMEM doorway endothelial cell (TEC, endomucin stained in blue, circled in white), TMEM doorway macrophage (TM, Iba1 stained in brown, circled in teal), and TMEM doorway tumor cell (TTC, Mena stained in pink, and circled in pink) and black arrows indicate where the cells are localized within the TMEM doorway. TMEM doorways were identified using automated analysis by VisioPharm identifying three adjacent immuno-histochemical stains. **C)** The sequential tissue sections after the IHC in (B) were stained with immunofluorescence (IF) with antibodies against endomucin (white), dextran (green), CSF-1 (red), and nuclear stain DAPI (blue). The two sequential sections from 1B and 1C were aligned and the same TMEM doorways were matched between the two sections, as indicated by the black circle in the IHC panel (1B) and white circle in the IF panels (1C) showing the same TMEM doorway after alignment. The left panels demonstrate the association of CSF-1 level in the TMEM doorway tumor cell (TTC) with TMEM doorway activity. The middle panel shows the extravascular signal for dextran as a green mask and the endomucin stain as a white mask, where thresholded, positive signal for these stains was converted into a binary mask. Active versus inactive TMEM doorways were distinguished by the presence of extravascular dextran staining non-overlapping with the endomucin stain, which indicates that the vessel had a TMEM doorway-associated vascular opening (TAVO). Scale bars=20μm. **D)** Immunofluorescence measurement of CSF-1 levels in TMEM doorway tumor cells in active and inactive TMEM doorways. TMEM doorways were identified in the IF-stained sections as described above. Active TMEM doorways were identified by the presence of extravascular dextran which is not present in inactive TMEM doorways (see panel C). Next, tumor cells were identified using the Mena positive cells at TMEM doorway (panel 1B, IHC stain). The level of CSF-1 in the TMEM doorway tumor cells was measured in both active and inactive TMEM doorways using Visiopharm. n= 12 mice. \*\*\*\**P* < 0.0001 analyzed by Student’s *t*-test.

### Tumor cells stimulate macrophage VEGF-A secretion without affecting its expression

Since CSF-1 is upregulated in the tumor cell at active TMEM doorways, and secretion of VEGF-A is known to be regulated by CSF-1 in other cell types (25, 26), we investigated the signaling interplay between the tumor cell and macrophage at the TMEM doorway. Using an ELISA on conditioned media, we confirmed that exogenous recombinant CSF-1 can stimulate macrophages to secrete VEGF-A **(Figure 2A)** and that tumor cells secrete CSF-1 **(Figure 2B)**. To examine the role of tumor cell secreted CSF-1 in regulating macrophage VEGF-A secretion, we first determined if TMEM doorway macrophages express the CSF-1 receptor (CSF-1R). We have previously identified TMEM doorway macrophages by their direct contact with a tumor cell and blood vessel (13), expression of CD68 and VEGF-A, and further characterized them as a CD206^+^/CD11b^+^/F4/80^+^/CD11c^-^ population (10). Using immunofluorescence staining of PyMT tumors for CSF-1R, macrophages (CD68^+^/CD206^+^/CD31^-^), and endothelial cells (CD31^+^) as described previously (10, 32), we found that greater than 90% of all CD68^+^ macrophages throughout the tumor tissue co-express CSF-1R and 97% of CD68^+^/CD206^+^ TMEM doorway macrophages (identified by simultaneous contact with tumor cells and endothelial cells and co-expression of CD68 and CD206 without CD31 expression) also express CSF-1R **(Figure 2C, D)**.

**Figure 2.**
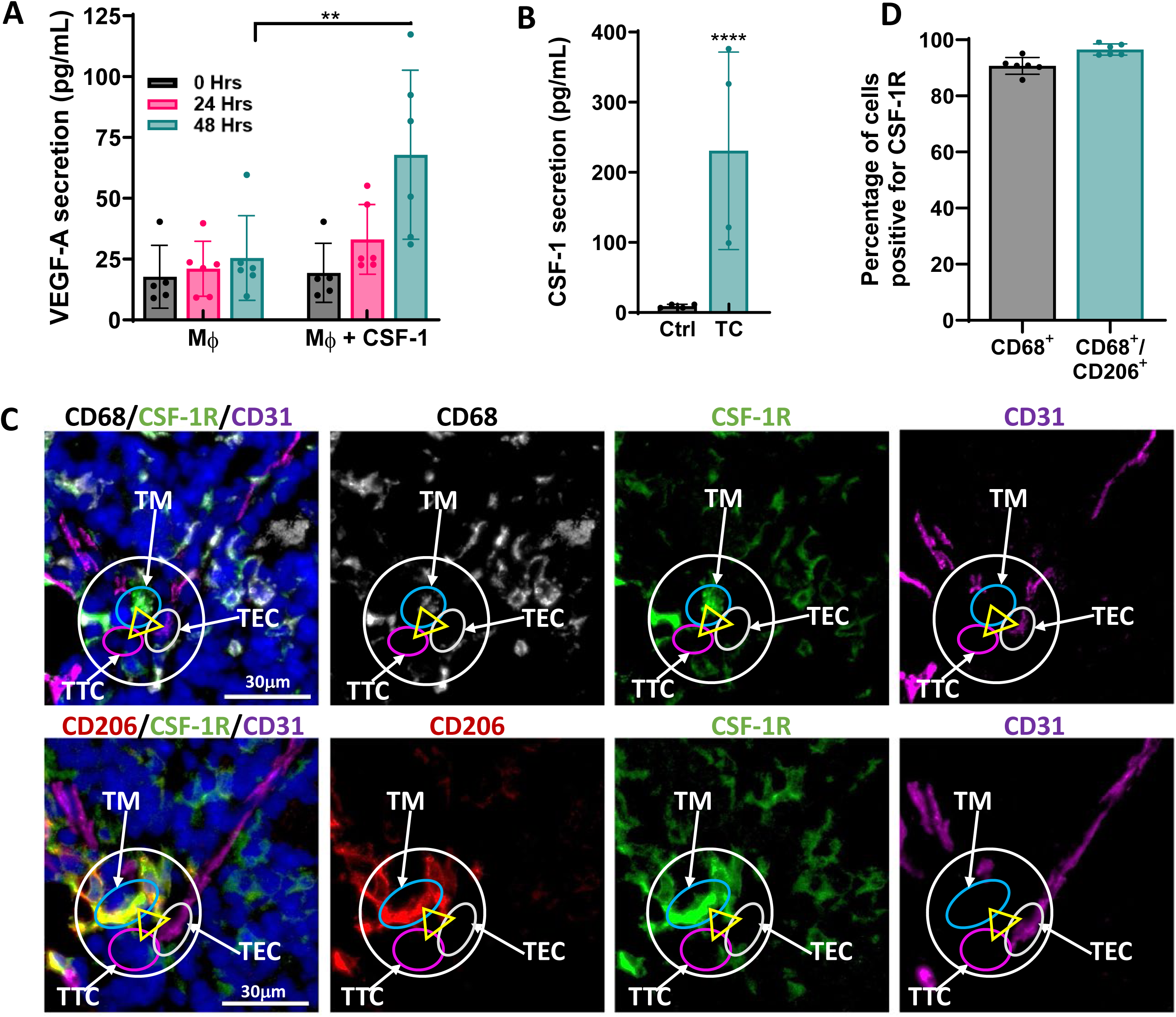
TMEM doorway macrophages secrete VEGF-A in response to CSF-1 and tumor cells secrete CSF-1. **A)** VEGF-A ELISA of conditioned media obtained from BAC1.2F5 macrophages (Mϕ) treated with or without CSF-1 (3000 units/mL) for the times indicated. VEGF-A is indicated as concentration (pg/ml) of secreted protein. n=3 individual experiments done in duplicate, **p<0.01, by two-way ANOVA. **B)** CSF-1 ELISA of conditioned media obtained from MDA-MB 231 tumor cells (TCs). TCs were cultured for 24 hours in serum-free media and the concentration of CSF-1 (pg/mL) secreted by the TCs measured in the tumor cell conditioned media by ELISA. Control is media not exposed to tumor cells but treated in the same way as the cells. n=3 individual experiments done in duplicate. ****p<0.0001 by Student’s *t-*test. **C)** Immunofluorescence staining of serial PyMT tumor sections stained for CSF-1R (green), vasculature (CD31, magenta), macrophages (top: CD68, white; bottom: CD206, red) and DAPI (blue). The TMEM doorway is indicated with the yellow arrow, with each point of the triangle identifying each cell in the doorway, as described in Figure 1 (10). TMEM doorway cells are indicated with white arrows: endothelial cell (TEC, white circle); TMEM doorway macrophage (TM, blue circle); and TMEM doorway tumor cell (TTC, pink circle). Scale bar 30 µm. **D)** Quantification of the percent of CD68^+^ TMEM doorway macrophages and CD68^+^/CD206^+^ macrophages which stain positively for CSF-1R (*n* = 6).

To determine if tumor cells can stimulate macrophage VEGF-A secretion, we used a human tumor cell - mouse macrophage co-culture system and quantified the amount of secreted VEGF-A by ELISA using an antibody that can only recognize mouse-derived VEGF-A, secreted by the macrophages. Co-culture of human MDA-MB-231 tumor cells with mouse BAC1.2F5 macrophages **(Figure 3A)** or culture of BAC1.2F5 macrophages with conditioned media collected from serum starved MDA-MB-231 tumor cells **(Figure 3B)**, resulted in increased secretion of VEGF-A by the mouse-derived macrophages.

**Figure 3.**
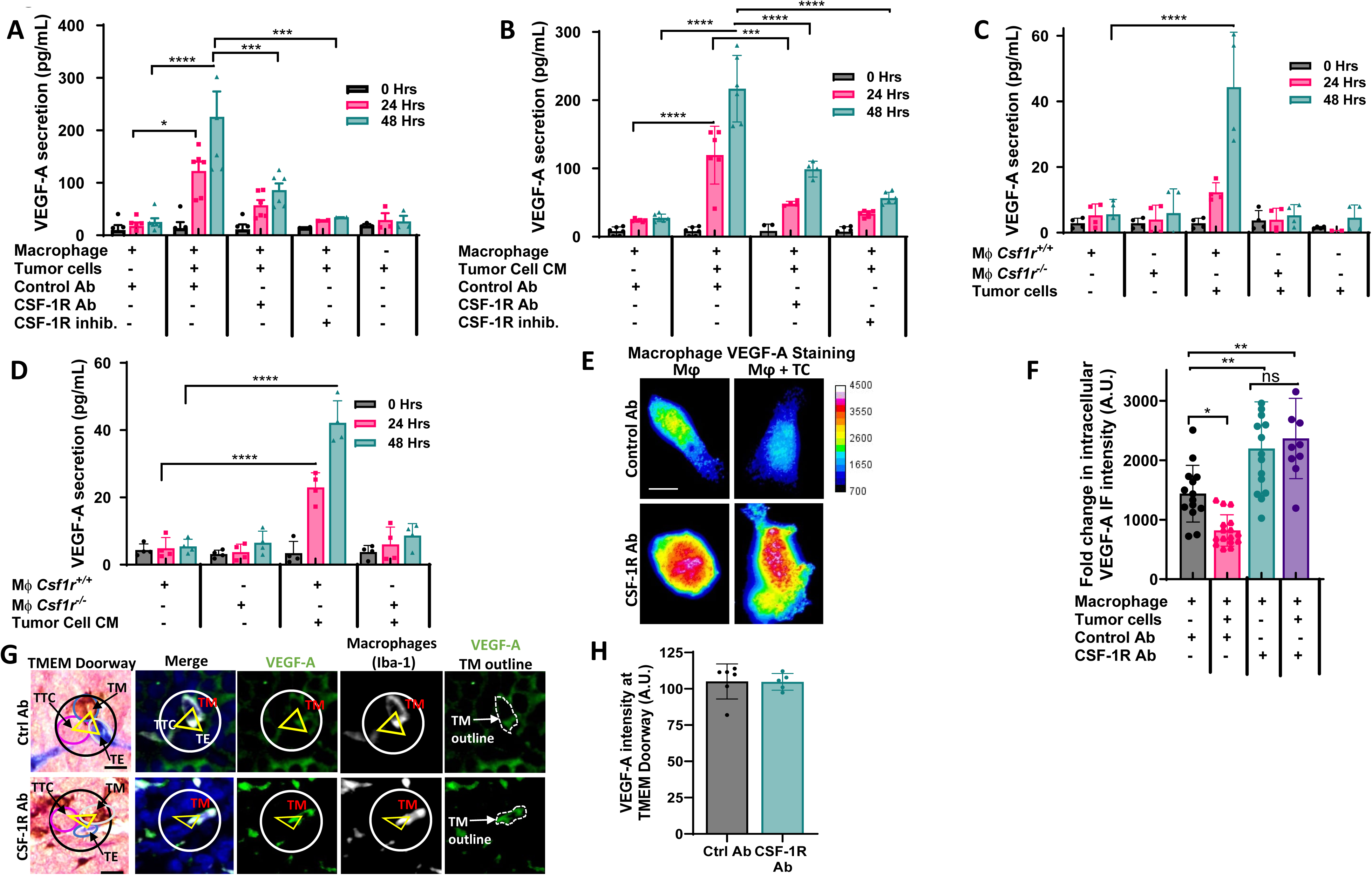
Tumor cell secreted CSF-1 increases CSF-1R dependent macrophage VEGF-A secretion. **A)** VEGF-A ELISA of conditioned media obtained from BAC1.2F5 macrophages co-cultured with MDA-MB-231 tumor cells treated with control antibody (ctrl Ab), CSF-1R blocking Ab or CSF-1R inhibitor (GW2580), denoted as concentration (pg/mL) of secreted protein. n=3 individual experiments performed in duplicate, *p<0.05, ***p<0.001, ****p<0.0001 analyzed by two-way ANOVA. **B)** ELISA determination of VEGF-A (pg/mL) in medium obtained from BAC1.2F5 macrophages cultured in medium conditioned by MDA-MB-231 tumor cells (Tumor Cell CM) and treated with control Ab, CSF-1R blocking Ab (CSF-1R Ab) or CSF-1R inhibitor (GW2580). n=3 individual experiments performed in duplicate, ***p<0.001, ****p<0.0001 analyzed by two-way ANOVA. **C)** ELISA determination of VEGF-A (pg/mL) in medium conditioned y bone marrow macrophage (BMMs) either expressing (*Csf1r^+/+^*) or lacking (*Csf1r^-/-^)* CSF-1R co-cultured with MDA-MB-231 tumor cells for the times indicated. n=3 individual experiments done in duplicate, ****p<0.0001 analyzed by two-way ANOVA. **D)** ELISA determination of VEGF-A (pg/mL) in medium from *Csf1r^+/+^* or *Csf1r^-/-^* BMMs cultured in MDA-MB-231 tumor cell conditioned medium (Tumor Cell CM) for the times indicated. n=3 individual experiments done in duplicate, ****p<0.0001 analyzed by two-way ANOVA. **E)** BAC1.2F5 macrophages, labelled with CellTracker^TM^ Green, were co-cultured with or without MDA-MB-231 tumor cells (TC), labelled with CellTracker^TM^ Red, and treated with either control or CSF-1R blocking antibodies for 48 hours. Cells were fixed, permeabilized and stained for VEGF-A. Macrophages were identified by the CellTracker Green label and only images of macrophages are shown. Heat map scale to right shows intensity of VEGF-A staining. Scale bar 5 µm. **F)** Quantitation of the VEGF-A fluorescence intensity in macrophages in (E). The amount of VEGF-A in the macrophage was quantified using ImageJ, as described in the methods section. n=3. *p<0.05, **p<0.01, analyzed by one-way ANOVA. **G, H)** Immuno-staining and quantification of VEGF-A intensity around TMEM doorways, obtained from PyMT mice treated with control antibody (ctrl Ab) or CSF-1R blocking Ab. Sequential tumor sections derived from PyMT tumors were stained by immunofluorescence (VEGF-A, Iba-1) and IHC (TMEM doorways-Mena, Iba-1, endomucin). TMEM doorways were identified as described in figure 1A. The circle in the IHC (black) and IF (white) panels show the same TMEM doorways obtained from the alignment of serial sections, and the three cells making up the TMEM doorway are pointed out with the yellow triangle in each panel. TMEM doorways are outlined in the circle with the endothelial cell (TE, blue stain, white circle), macrophage (TM, brown stain, teal circle) and TMEM doorway tumor cell (TTC, pink stain, pink circle) indicated with arrows. Next, the two sequential sections were aligned and TMEM doorway was identified in the IF-stained section. The immunofluorescence intensity of VEGF-A expression (green stain) in the identified TMEM doorway ROIs was measured and plotted here. N=analysis of images obtained from tissue sections from individual 11 mice, Scale bar 20 µm, not significant, analyzed by Student’s *t*-test.

To ascertain the role of tumor cell secreted CSF-1 in stimulating macrophage VEGF-A secretion, we used the same co-culture system of MDA-MB-231 tumor cells and BAC1.2F5 macrophages as above. We inhibited CSF-1R signaling in macrophages with either a CSF-1R neutralizing antibody (CSF-1R Ab), or an inhibitor which prevents downstream signaling of the CSF-1R (CSF-1R inhibitor (GW2580)). We found that either method reduced macrophage VEGF-A secretion **(Figure 3A)**. Likewise, inhibition of CSF-1R in macrophages cultured with medium conditioned by the tumor cells led to a reduction in the amount of secreted VEGF-A **(Figure 3B)**. Similar results were obtained using mouse bone marrow derived primary macrophages (BMMs) **(Supplementary Figure 1A)**. Co-culturing a different breast cancer cell line, 4T1, which also secretes CSF-1 **(Supplementary Figure 1B)**, with BAC1.2F5 macrophages also caused an increase in VEGF-A secretion by macrophages **(Supplementary Figure 1C).** In these additional models, the increase in macrophage secretion of VEGF-A in response to tumor cell CSF-1 was again suppressed through addition of CSF-1R inhibitors or blocking antibodies **(Supplemental Figure 1A, 1C)**. Next, we used immortalized bone marrow macrophages (BMMs) isolated from *Csf1r^+/+^* or *Csf1r^-/-^* mice which either express (*Csf1r^+/+^*) or lack the expression of CSF-1R (C*sf1r^-/-^)* (33) to determine the role of the CSF-1R in regulating VEGF-A secretion by macrophages. Compared to the *Csf1r^+/+^* BMMs, which secreted VEGF-A in both the co-culture system and when cultured in tumor cell conditioned media **(Figure 3C, D)**, C*sf1r^-/-^* BMMs did not significantly secrete VEGF-A, in either a co-culture system with MDA-MB-231 tumor cells, or when cultured with medium conditioned by the tumor cells **(Figure 3C, D)**.

We next evaluated if co-culture of tumor cells with macrophages leads to increased expression or secretion of VEGF-A from macrophages, or both. We co-cultured MDA-MB-231 tumor cells with BAC1.2F5 macrophages labeled with CellTracker^TM^ Green to distinguish them from cancer cells, and then stained the cells for VEGF-A. Macrophages that were co-cultured with tumor cells displayed decreased intracellular VEGF-A staining compared to macrophages cultured alone **(Figure 3E, F)**. This result, combined with the VEGF-A ELISA showing that macrophages and tumor cells cultured in the same method lead to increased VEGF-A in the conditioned media (Figure 3A), suggests that the tumor cells induce macrophages to secrete VEGF-A. In addition, inhibition of CSF-1R signaling (which prevented VEGF-A secretion in Figure 3A) increased the intracellular macrophage VEGF-A using the same co-culture system **(Figure 3E, F)**, indicating that the CSF-1/CSF-1R signaling between macrophages and tumor cells induces the release of intracellular VEGF-A from macrophages. In our second model using 4T1 tumor cells, we again found a decrease in intracellular VEGF-A staining in the macrophages co-cultured with tumor cells compared to macrophages cultured without tumor cells **(Supplementary Figure 1D).** The increase in macrophage VEGF-A secretion and the decrease in the intracellular VEGF-A protein levels in the macrophages were abrogated with CSF-1R neutralizing antibody as well as with the small molecule CSF-1R inhibitor, consistent to what was observed with MDA-MB-231 cells in **Figure 3**.

To determine if the enhancement of VEGF-A secreted by the macrophages in response to co-culture with tumor cells could also be due to increased VEGF-A mRNA production, we co-cultured MDA-MB-231 tumor cells with BAC1.2F5 macrophages and determined the relative levels of VEGF-A mRNA by qPCR. We found that co-culture of tumor cells and macrophages did not affect the VEGF-A mRNA levels in macrophages **(Supplementary Figure 2)**, indicating that the increase in VEGF-A protein found in the conditioned medium was due to increased secretion of stored intracellular VEGF-A in the macrophages, not also caused by an increase in VEGF-A mRNA expression.

To determine the impact of CSF-1R inhibition on the levels of VEGF-A expression *in vivo*, we treated mice with either a control or a CSF-1R inhibitory antibody and looked at the total level of VEGF-A (both intra- and extracellular) within TMEM doorways. We found that CSF-1R inhibition does not affect the steady state expression or stability of VEGF-A levels in the TMEM doorway. **(Figure 3G, H)**. Both findings are consistent with the observation that tumor cell secreted CSF-1 does not upregulate VEGF-A expression in macrophages **(Supplementary Figure 2)**, but rather controls VEGF-A secretion from macrophages, which is retained within the TMEM doorway. Altogether, these results indicate that tumor cells stimulate macrophages to release VEGF-A but do not induce its production within macrophages.

### Inhibition of CSF-1R reduces TMEM doorway-associated vascular opening and trans-endothelial migration

As CSF-1R signaling regulates macrophage VEGF-A secretion, and macrophage-secreted VEGF-A is required for vascular opening at TMEM doorways (10), we examined the role of the CSF-1R in mediating vascular opening and tumor cell intravasation at TMEM doorways *in vivo,* as defined in **Figure 1** and previously (6). We injected tumor bearing PyMT mice *i.v.* with CSF-1R blocking antibodies or IgG control antibodies, four hours prior to sacrifice, and injected high molecular weight dextran (155 kDa), one hour before sacrifice. We stained the tumor tissue for CD31, TMR-Dextran, and ZO-1 to examine the effects of CSF-1R inhibition on TMEM doorway activity and vascular junctional integrity (**Figure 4A).** Acute inhibition of CSF-1R reduced extravascular dextran (red) at TMEM doorways, indicating that TMEM doorways were less active when CSF-1 signaling is inhibited **(Figure 4B)**. Vascular ZO-1, a component of endothelial tight junctions that maintains vascular junction integrity, increased with inhibition of CSF-1R suggesting that the endothelial junctions were less permeable upon CSF-1R inhibition **(Figure 4C)**. To determine whether blockade of CSF-1 signaling and subsequent inhibition of TMEM doorway opening prevented tumor cells from entering the vasculature, we collected the blood of the mice during sacrifice and measured the number of circulating tumor cells (CTCs). The number of CTCs were significantly reduced upon CSF-1R inhibition compared to control treated mice **(Figure 4D)**. These results indicate that CSF-1R signaling is not only involved in tumor cell and macrophage streaming migration, as reported before (8), but also in signaling between tumor cells, macrophages, and endothelial cells, creating intravasation-associated vascular openings and allowing intravasation of tumor cells, leading to CTCs.

**Figure 4.**
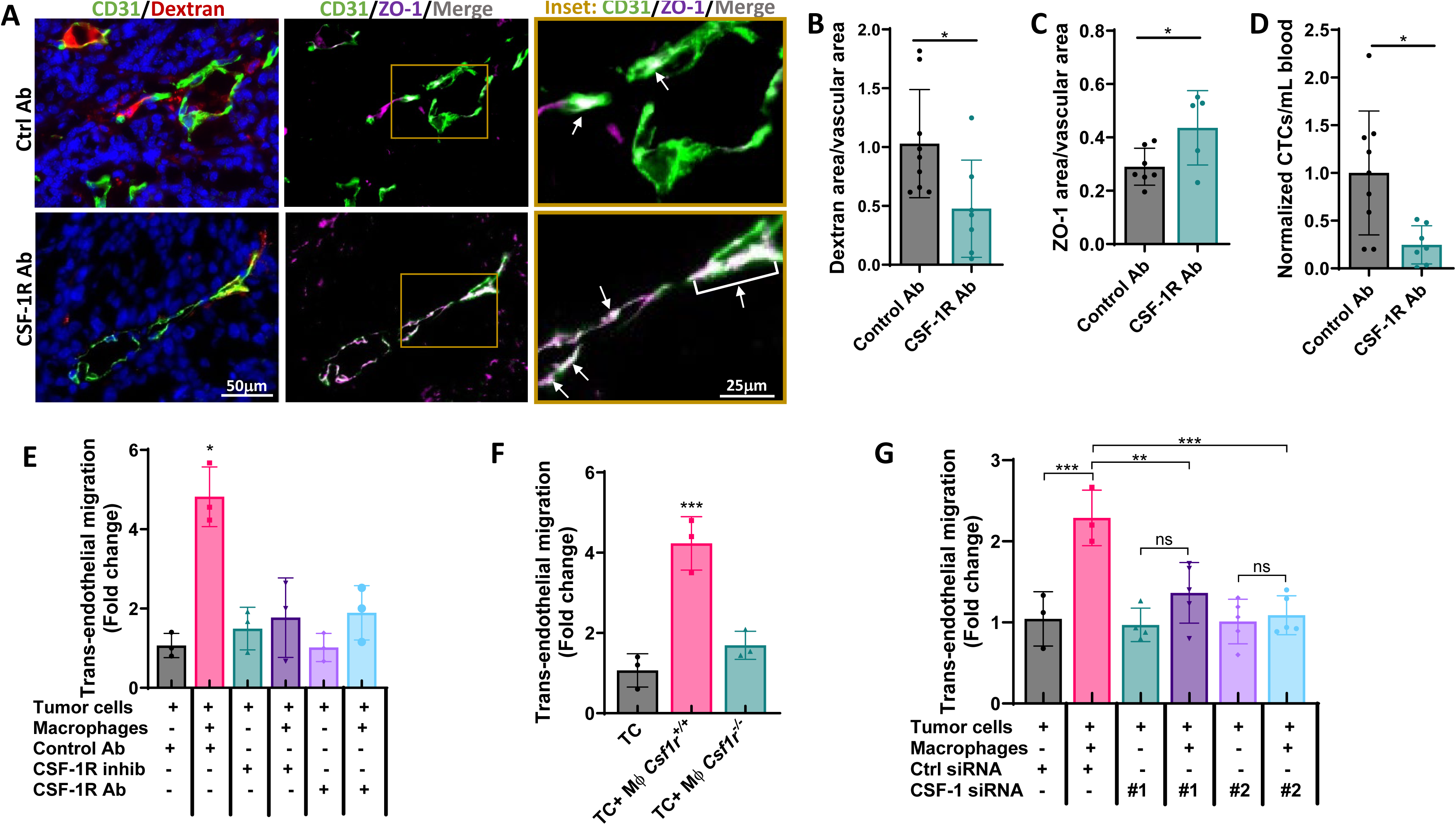
Inhibition of CSF-1R signaling reduces TAVO and trans-endothelial migration. **A)** IF staining of tumor tissue from mice treated with control Ab or CSF-1R Ab were stained for CD31 (green), TMR-dextran (red), and ZO-1 (magenta). Right panel shows increased magnification of yellow box from middle panel to demonstrate overlap between CD31 and ZO-1 stains (merge/overlap indicated with white signal, and pointed out with white arrows). **B)** Quantification of extravascular 155 kDa dextran-TMR. *p<0.05. **C**) Vascular ZO-1 *p<0.05, and **D**) circulating tumor cells for mice treated with control Ab or CSF-1R blocking Ab in A). *p<0.05, analyzed by Student’s *t*-test. (B, C, D, *n* = 8) **E)** Quantitation of subluminal to luminal trans-endothelial migration (iTEM activity, which is a measure of number of intravasating cells in the intravasation direction) of MDA-MB-231 cells plated on the confluent and sealed endothelium either alone or with BAC1.2F5 macrophages. Tumor cells were treated with control antibody (ctrl Ab & DMSO), CSF-1R blocking Ab (CSF-1R Ab) or CSF-1R inhibitor (GW2580). Tumor cells were labelled with CellTracker^TM^ green which allowed us to identify and quantify the cells crossing. The number of MDA-MB-231 cells which crossed the endothelium were imaged using a confocal microscope and quantified using ImageJ software. n=3, *p<0.05 analyzed by one-way ANOVA. **F)** Quantitation of iTEM activity of MDA-MB-231 cells plated on the endothelium alone or with BMMs expressing or lacking CSF-1R expression. Tumor cells were labelled with CellTracker^TM^ green which allowed us to identify and quantify the cells crossing. The number of MDA-MB-231 cells which crossed the endothelium were imaged using a confocal microscope and quantified using ImageJ software. N=3, ***p< 0.001 analyzed by one-way ANOVA. **G)** Quantitation of iTEM activity of MDA-MB-231 cells transfected with either control siRNA or two different siRNAs targeting CSF-1 (#1, #2). Transfected cells were plated on the endothelium either alone or with BAC1.2F5 macrophages. Tumor cells were labelled with CellTracker^TM^ green which allowed us to identify and quantify the cells crossing. The number of MDA-MB-231 cells which crossed the endothelium were imaged using a confocal microscope and quantified using ImageJ software. n=3, **p<0.01, ***p< 0.001 analyzed by one-way ANOVA.

To further examine the effects of blocking CSF-1R signaling on macrophage-mediated transendothelial migration of tumor cells, we used our previously established *in vitro* trans-endothelial migration (iTEM) assay (34). The iTEM assay mimics the conditions seen at the TMEM doorway during intravasation (10, 32) and allows us to quantify the number of tumor cells that cross the endothelium. We have previously shown that breast tumor cells seeded alone atop a matrigel matrix and confluent endothelial monolayer cross through the endothelial cell barrier poorly, but transendothelial migration is enhanced in the presence of macrophages (34, 35). As shown before, tumor cells exhibited a basal level of trans-endothelial migration, which was significantly increased in the presence of macrophages (**Figure 4E**). However, there was no enhancement of tumor cell trans-endothelial migration in presence of macrophages when CSF-1R signaling was inhibited by either a CSF-1R blocking antibody or a small molecule inhibitor of CSF-1R (**Figure 4E**).

To further establish that CSF-1R signaling between tumor cells and macrophages was important for enhancing tumor cell migration across the endothelial cell layer, we used BMMs lacking CSF-1R (*Csf1r^-/-^*) in the iTEM assay. We observed no increase in the transmigration of tumor cells seeded with *Csf1r^-/-^* macrophages compared to tumor cells cultured without macrophages; however, tumor cell transendothelial migration was enhanced when seeded with macrophages expressing the CSF-1R (*Csf1r^+/+^*) **(Figure 4F)**. Similarly, to establish that CSF-1 secreted by tumor cells specifically is responsible for the increased trans-endothelial migration, and not the other TMEM doorway cells present in the iTEM assay, we knocked down CSF-1 in MDA-MB-231 cells using siRNA (**Supplementary Figure 3**) and used these tumor cells in the iTEM assay. We found that unlike tumor cells expressing CSF-1, tumor cells with reduced CSF-1 expression did not support increased trans-endothelial migration in the presence of macrophages (**Figure 4G**). Thus, inhibition of CSF-1R signaling blocked the ability of macrophages to enhance tumor cell trans-endothelial migration.

## Discussion

It has been previously shown that TMEM doorway associated vascular opening (TAVO) requires vascular endothelial growth factor-A (VEGF-A) produced by the TMEM doorway macrophage. In particular, a TIE2^hi^/VEGF-A^hi^ TMEM doorway-associated macrophage interacts with its associated endothelial cell through secretion of VEGF-A to mediate blood vessel opening by disrupting endothelial cell adherens and tight junctions (10). We have demonstrated, using intravital imaging, that the vascular opening in tumors is an acute, localized, and transient event, and only occurs at TMEM doorways, as opposed to VEGF-A injected into the vasculature, which caused general and continuous leakage of blood contents from all blood vessels (10). We have further shown that these TAVO events occur simultaneously with tumor cell intravasation, and are required for tumor cell intravasation. When VEGF-A is knocked out specifically in macrophages, TAVO events are blocked, as is tumor cell intravasation (10). These observations are consistent with and provide a mechanism for the finding that the number of TMEM doorways measured by multiplex immunohistochemistry in formalin-fixed paraffin-embedded breast cancer sections is a clinically validated prognostic indicator of distant metastasis in breast cancer patients (12–14). However, the signal provided by CSF-1 that triggers the secretion of VEGF-A by the TMEM doorway macrophage, the most critical step for TMEM doorway opening, has remained unknown, until now.

CSF-1 drives the differentiation of myeloid precursors to macrophages and is essential for survival of macrophages (36–38). CSF-1/CSF-1R signaling is important for tissue homeostasis, repair, and inflammation and acts as a chemoattractant for the recruitment, production, and survival of tumor associated macrophages (TAMs) to the tumor microenvironment (39, 40). TAMs promote tumor progression through stimulation of angiogenesis, secretion of IL-10, TGFβ, and VEGF-A, and driving of tumor cell migration, invasion, and metastasis (41–43). As we show here, VEGF-A secretion from TMEM doorway macrophages can be triggered by tumor cell derived-CSF-1 binding to the CSF-1R on the macrophage (Figures 2 and 3). Given that CSF-1 signaling is known to drive invasiveness and increase metastasis, unsurprisingly, increased expression of CSF-1 is correlated with poor prognosis in ovarian, breast, and prostate cancer patients (44–49). There is evidence of a paracrine interaction between tumor cells and macrophages, which facilitates tumor cell migration toward blood vessels in breast cancer (8, 29). This interaction is driven by a CSF-1/EGF paracrine loop, wherein tumor cells secrete CSF-1 (which attracts CSF-1R expressing macrophages), while macrophages in turn secrete EGF (which attracts EGFR-expressing tumor cells). The tumor cells respond to EGF-secreting macrophages by migrating towards them and secreting more CSF-1, therefore generating a paracrine loop (8, 27, 29, 50). These signaling events create a chemotactic axis in which the CSF-1/EGF signals become amplified, allowing the tumor cells and macrophages to remain in close proximity with each other as they migrate together as cell pairs (a process called streaming (8, 27, 29, 36)). Once the tumor cells within the loop encounter a Hepatic Growth Factor (HGF) signal, which is found in a gradient in the tumor microenvironment with the highest concentration near blood vessels (7), the tumor cell-macrophage pairs start migrating toward the blood vessels along the HGF gradient, adding directionality to the streaming cell pairs **(Figure 5A)**. Once at the blood vessels, the streaming tumor cells can intravasate through active TMEM doorways, following a TAVO event (7, 10, 11). The genetic or pharmacological inhibition of EGF, CSF-1, or the HGF receptor (c-Met) significantly impairs streaming of both tumor cells and macrophages and blocks tumor cell dissemination in *in vivo* breast cancer metastasis models (7, 8, 29, 50), confirming the critical role of the paracrine CSF-1/EGF loop between tumor cells and macrophages, as well as the HGF gradient, in metastasis.

**Figure 5.**
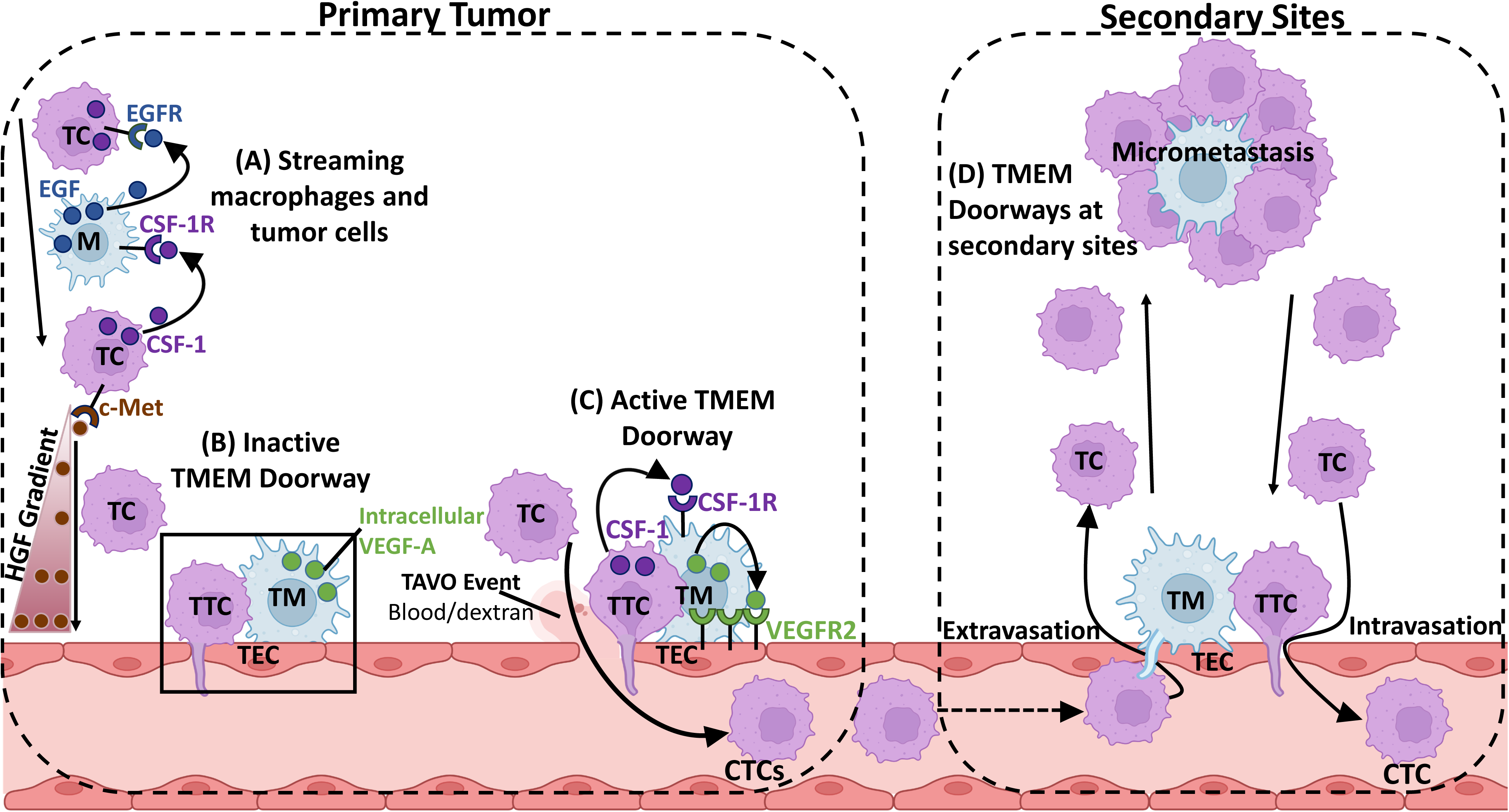
Model of CSF-1-induced TMEM doorway opening and release of CTCs into the blood stream. **A)** In the primary tumor, macrophages and tumor cells exhibit a CSF-1/EGF paracrine interaction between tumor cells and macrophages which facilitates tumor cell migration toward blood vessels in breast cancer. Tumor cells secrete CSF-1 which attracts CSF-1R expressing macrophages, which in turn secrete EGF, increasing the migration of EGFR-expressing tumor cells. The tumor cells within the loop encounter an HGF signal found in a gradient in the tumor microenvironment with the highest concentration near blood vessels. The tumor cells migrate toward the blood vessels along the HGF gradient, adding directionality to the migration of the tumor cell-macrophage paracrine loop engaged partners. **B)** In an inactive TMEM doorway (box), with the absence of any CSF-1 secreted by the TMEM doorway tumor cell (TTC), the VEGF-A remains within the TMEM doorway macrophage (TM). The blood vessels, including the TMEM doorway endothelial cell (TEC) within the tumor remain sealed and do not allow tumor cells to intravasate through TMEM doorways. **C)** In contrast, in an active TMEM doorway, CSF-1 is secreted by the TMEM doorway tumor cell and binds to macrophage CSF-1R, stimulating the TMEM doorway macrophage to secrete its VEGF-A. VEGF-A causes vascular opening (TAVO event) and allows tumor cells to intravasate through the TMEM doorway, creating CTCs, and dextran or blood to leak out of the vessel. **D)** These circulating tumor cells (CTCs) travel through the vasculature to secondary sites where TMEM doorways are also found in metastatic foci in lymph nodes and lungs. Thin membranous connections between macrophages and tumor cells stretch from the macrophage on extravascular side of the blood vessel through the endothelial junctions and interact with CTCs which could facilitate CTC extravasation at secondary sites. TMEM doorways in metastases could also be the sites were tumor cells re-intravasate and then seed tertiary metastases. Figure created with BioRender.com.

Here we demonstrate that while the TMEM doorway tumor cell and macrophage remain in direct and stable contact on the blood vessel (10, 13), the CSF-1 signaling persists, resulting in localized macrophage secretion of VEGF-A. We found that TMEM doorway tumor cells secrete CSF-1 which binds to the CSF-1R on TMEM doorway macrophages **(Figure 5B)**. We further show that this event triggers localized secretion of VEGF-A by TMEM doorway macrophages and leads to downstream TAVO events, where acute vascular opening within the endothelium allows tumor cells to intravasate through the TMEM doorway. Previously, we have visualized and characterized that the resulting CTCs can form distant metastases **(Figure 5C, D)** (10, 31, 51). In this study, we demonstrate that inhibition of CSF-1R *in vitro* by either knocking out CSF-1R in macrophages or adding a blocking antibody decreased macrophage release of VEGF-A, and transendothelial migration of tumor cells, an *in vitro* measure of intravasation capability. *In vivo*, acute treatment of tumor bearing mice with a CSF-1R blocking antibody significantly increased vascular tight junction stability, decreased TMEM doorway activity and TAVO events, as well as formation of CTCs. We did not detect a difference in VEGF-A expression around the TMEM doorway when a CSF-1R blocking antibody was used *in vivo* (Figure 3G, 3H). Given that this is a relatively short-term experiment, sacrificing mice four hours after treatment, and the TAVO is a rare event, we expect that detecting a change in VEGF-A expression within the TMEM doorways is not possible in this limited timeframe. Further, we have found that CSF-1 signaling does not increase expression of VEGF-A in the TMEM doorway macrophage, but rather triggers its local secretion, where VEGF-A binds to and signals to the nearby TMEM doorway endothelial cell, while remaining within the TMEM doorway.

The importance of this study beyond the primary tumor is emphasized by the finding that TMEM doorways are found not only in invasive ductal breast carcinomas, but also in metastatic foci in lymph nodes (18, 19) and lungs (6). We and others have found that secondary metastases can seed tertiary metastases (1–4), and the presence of TMEM doorways at secondary sites may perpetuate metastatic dissemination **(Figure 5D)** even after removal of the primary tumor. Importantly, blocking the function of TMEM doorways could not only prevent the dissemination of cancer cells from primary tumors, but also prevent re-dissemination of tumor cells from metastatic sites, which may occur after the resection of primary tumors (10, 11, 52, 53). Therefore, even if the tumors are highly metastatic, we may be able to increase patient survival or quality of life, or both, by blocking TMEM doorway function systemically, which would lessen overall metastatic burden. Inhibition of both Tie2 and CSF-1 signaling to prevent TAVO events (10, 31), along with standard of care therapies, might be an effective way to prevent re-dissemination from both primary and metastatic tumor sites, and allow for better patient survival.

Recent preclinical studies targeting CSF-1/CSF-1R have yielded promising results in several tumor models. A monoclonal antibody targeting CSF-1R depleted TAMs and increased the CD8^+^/CD4^+^ T cell ratio, leading to decreased primary tumor burden and decreased metastasis in mouse models of colorectal cancer and fibrosarcoma (54). Another recent study investigating breast and colon carcinoma models found that CSF-1/CSF-1R signaling inhibition stimulated the expansion of neo-epitope specific T cells and promoted the development of an immune-permissive TME (55). So far, the FDA has approved one CSF-1R inhibitor for patients with tenosynovial giant cell tumors, a tumor type characterized by chromosomal translocations of the CSF-1 gene, leading to aberrant CSF-1 expression (56, 57). Despite the promising preclinical studies using CSF-1/CSF-1R inhibitors in combination with chemotherapy, immunotherapy, or radiotherapy, phase II clinical trials have not yielded the same anti-tumor efficacies (58–64). In patients with advanced triple negative breast cancer, the combination of a CSF-1 monoclonal antibody with gemcitabine and carboplatin showed comparable anti-tumor activity compared to gemcitabine and carboplatin alone (65). In another phase Ib/II study, patients with advanced pancreatic ductal carcinoma, colorectal cancer, or non-small cell lung cancer were treated with a CSF-1R monoclonal antibody plus anti-PD-1 therapy; however, minimal anti-tumor activity was observed (62). In these studies, investigators were expecting to observe an enhancement in intratumoral infiltration of CD8^+^/CD4^+^ T cells and TAM depletion, given the role that CSF-1 signaling plays in creating an immune suppressive environment and promoting recruitment, survival and differentiation of macrophages. Taking into consideration our findings that CSF-1 signaling is critical for the recruitment of tumor cell-macrophage pairs to the blood vessels in the primary tumor (8, 27, 29, 50), the transient opening of the blood vessels at TMEM doorways (Figure 4B), and intravasation of tumor cells at TMEM doorways (Figure 4D), we expect that treating patients with less advanced disease would be efficacious in preventing metastatic disease or reducing metastatic burden, though not necessarily in preventing primary tumor growth. Additionally, given our findings that TMEM doorway scores are predictive of distant metastasis in subsets of breast cancer patients (12–14), we contend that selecting the breast cancer patients meeting the criteria for predictive metastatic disease for the clinical trials would show increased success in improving patient outcomes. Our study provides evidence that the mechanism through which CSF-1R inhibition can prevent metastasis is not only by macrophage depletion, but also by blocking CSF-1/CSF-1R paracrine signaling between tumor cells and macrophages leading to inhibition in tumor cell intravasation in the primary tumor.

We have previously shown in mice that transient, short-term blockade of VEGF-A expression in macrophages is effective in inhibiting TMEM doorway activity, TAVO events, and formation of CTCs (10). However, if patients are treated with anti-angiogenic therapies targeting VEGF-A (e.g., bevacizumab and other VEGF-A inhibitors) they eventually develop therapeutic resistance (66). One mechanism of tumor resistance or recurrence following anti-angiogenic therapy in patients is attributed to the increased recruitment and infiltration of Tie2-expressing macrophages into the tumor in response to apoptosis, necrosis, and hypoxia after vascular regression (66). Additionally, the ligand for Tie2, Ang2, is upregulated in response to hypoxic conditions. Tie2-Ang2 signaling can function in a similar manner to VEGF-A signaling to promote tumor angiogenesis, bypassing VEGF-A inhibition, and even being enhanced by such inhibition. As such, Tie2 is an attractive pharmacological target for the suppression of tumor angiogenesis and tumor cell dissemination. We have found that TMEM doorway macrophages are also characterized by expression of Tie2 (10) and that a selective inhibitor of Tie2, rebastinib, inhibits TMEM doorway activity, formation of CTCs, and metastasis in several mouse models of metastasis and patient derived xenografts (32). The mechanism of how Tie2 contributes to signaling between the TMEM doorway cells to mediate the TAVO events is still unclear and warrants further investigation. Uncovering the complete mechanism of TMEM doorway opening in primary and secondary sites is critical. We expect that inhibiting TAVO events, potentially by targeting both CSF-1 and Tie2 signaling, will improve our ability to increase the survival of both patients with metastatic disease and patients who are likely to develop metastatic disease.

## Acknowledgments

This work was supported by the NCI grants (R01 CA255153, PPG CA257885, R01 CA240646, F32 CA243350), SIG OD019961, the Gruss-Lipper Biophotonics Center, the Integrated Imaging Program, the Integrated Imaging Program for Cancer Research, The Evelyn Gruss-Lipper Charitable Foundation, and The Helen & Irving Spatz Family Foundation. We would like to thank the Analytical Imaging Facility at Albert Einstein College of Medicine for imaging support.

## Author contributions

Conceptualization: CRS, ASH, MHO, JSC

Methodology: CRS, ASH, XY,

Formal Analysis: CRS, CLD, XY, VPS, DE, MHO

Software: XY, DE

Investigation: CRS, CLD, XC, YL, ASH, YW, VPS

Resources: ERS, DC, DE

Writing: CRS, CLD, ASH, ERS, MHO, JSC

Funding Acquisition: CLD, DE, MHO, JSC

Supervision: ERS, JCM, DE, MHO, JSC

## Data availability statement

Data sharing is not applicable to this article as no datasets were generated or analyzed during the current study.

