## Supplemental Figures and Legends for "Signaling events at TMEM doorways provide potential targets for inhibiting breast cancer dissemination"

**Supplemental Figure Legends:**

**Supplementary Fig. 1 CSF-1 is required for bone marrow derived macrophage VEGF-A secretion and tumor cell secreted CSF-1 increases macrophage VEGF-A secretion. A)** ELISA determination of VEGF-A (pg/mL) in the medium conditioned by macrophages (BMMs) co-cultured with or without tumor cells (MDA-MB 231), with conditioned medium by MDA-MB-231 tumor cells (tumor cell CM), or in presence of CSF-1R small molecule inhibitor (GW2580, CSF-1R inhib.) or vehicle (DMSO). n=3 individual experiments performed in duplicate, ****$p<0.0001$ by two-way ANOVA. **B)** CSF-1 ELISA determination of conditioned media obtained from 4T1 tumor cells. 4T1 tumor cells were cultured for 24 hours in serum-free media and the concentration of CSF-1 (pg/mL) secreted by the 4T1 cells was measured in the tumor cell conditioned media by ELISA. Control is media not exposed to tumor cells but treated in the same way as the cells. n=3, ***$p<0.001$, analyzed by Student's *t*-test. **C)** ELISA determination of VEGF-A concentration (pg/mL) in medium conditioned by macrophages (BAC1.2F5) co-cultured with 4T1 tumor cells treated with control antibody (Ctrl Ab), CSF-1R blocking Ab (CSF-1R Ab), CSF-1R inhibitor (GW2580, CSF-1R inhib.) or DMSO control. n=2 individual experiments done in duplicate, *$p<0.05$, ***$p<0.001$, ****$p<0.0001$ analyzed by two-way ANOVA. **D)** Tumor cells decrease intra-cellular macrophage VEGF-A. Quantification of the immunofluorescence staining intensity of VEGF-A in macrophages (labelled with CellTracker™ Green) cultured with or without tumor cells 4T1 (labelled with CellTracker™ Red) and treated with either ctrl Ab or CSF-1R blocking Ab. Macrophages were labeled with CellTracker™ Green to distinguish them from tumor cells. The amount of VEGF-A in the macrophage was quantified using ImageJ. n=3. ****$p<0.0001$ analyzed by one-way ANOVA.

**Supplementary Figure 2: CSF-1 secreted by tumor cells does not increase macrophage VEGF-A mRNA levels.** Fold change VEGF-A mRNA expression, determined by qPCR, in macrophages (BAC1.2F5) co-cultured with tumor cells (MDA-MB-231) compared to macrophages cultured alone, without tumor cells. Murine VEGF-A mRNA expression was normalized to murine GAPDH mRNA expression, as the endogenous control. n=3 individual experiments performed in triplicate, non-significant (ns), analyzed by Student's *t*-test.

**Supplementary Figure 3: CSF-1 knock-down in MDA-MB-231 cells.** Fold change CSF-1 mRNA expression, determined by qPCR, in MDA-MB-231 cells transfected using Lipofectamine 2000 with ctrl siRNA or 3 different CSF-1 targeting siRNAs (#1, #2, #3) for 48hrs. Ctrl siRNA CSF-1 expression was set to one and all other treatments were set relative to this control. CSF-1 mRNA expression was normalized to GAPDH mRNA expression, as the endogenous control, in all treatment groups. n=3 individual experiments performed in triplicate, ***p<0.001, ****p<0.0001 analyzed by one-way ANOVA.

### Supplemental Figure 1

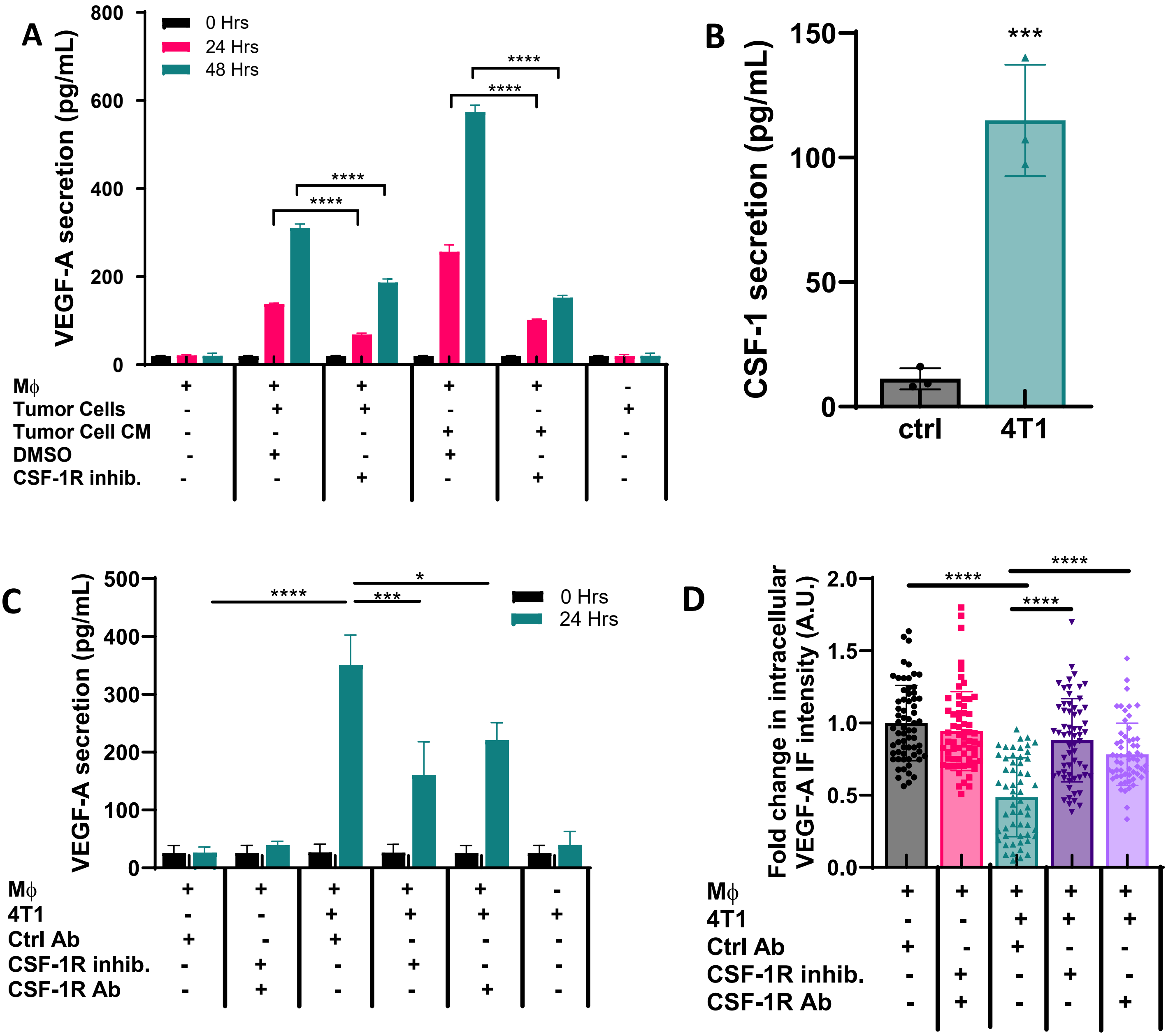

**Supplemental Figure 2**

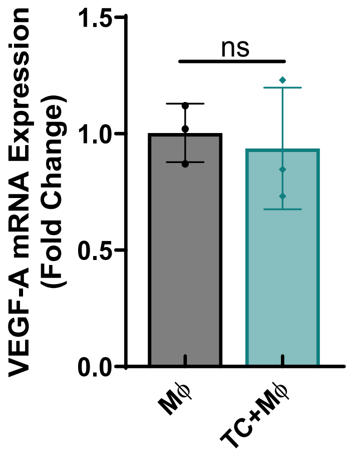

**Supplemental Figure 3**

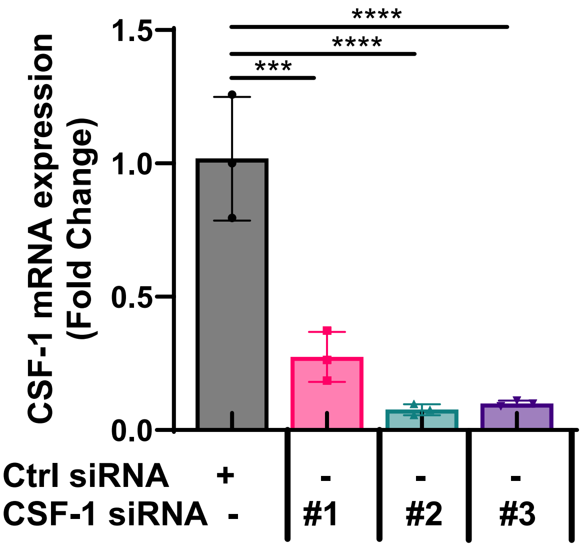
